# FGF9 and FGF10 use distinct signaling pathways to direct lung epithelial specification and branching

**DOI:** 10.1101/873109

**Authors:** Yongjun Yin, David M. Ornitz

## Abstract

Fibroblast Growth Factors (FGFs) 9 and 10 are essential during the pseudoglandular stage of lung development. Mesothelial produced FGF9 is principally responsible for mesenchymal growth, whereas epithelial produced FGF9 and mesenchymal produced FGF10 guide lung epithelial development, and loss of either of these ligands affects epithelial branching. Because FGF9 and FGF10 activate distinct FGF receptors (FGFRs), we hypothesized that they would control distinct developmental mechanisms. Here, we show that FGF9 signaled through epithelial FGF receptor 3 (FGFR3) to directly promote distal epithelial fate specification and inhibit epithelial differentiation. By contrast, FGF10 signaled through epithelial FGFR2b to promote epithelial proliferation and differentiation. Furthermore, FGF9-FGFR3 signaling functionally opposed FGF10-FGFR2b signaling, and FGFR3 preferentially used downstream PI3K pathways, whereas FGFR2b relied on downstream RAS-MAPK pathways. These data demonstrate that within lung epithelial cells, different FGFRs function independently; they bind receptor-specific ligands and direct unique developmental functions through activation of distinct downstream signaling pathways.

## Introduction

Mouse lung development begins with the emergence of two buds from the foregut at embryonic day 9.5 (E9.5). Between E9.5 and E16.5, the pseudoglandular stage of development forms the primary airway branching pattern of the lung. This is followed by the canalicular (E16.5-17.5), saccular (E18.5 to postnatal day 5 (P5)), and finally alveolar (P5-P21) stages that progressively form a functional adult lung (*1*). The lung bud contains three primary cell types: epithelium, mesenchyme, and mesothelium. The innermost layer, the epithelium, gives rise to the branched network of ducts that will conduct air to the alveoli, the distal gas-exchange units of the lung. Mesenchyme surrounding the epithelial ducts will form the vascular and stromal components of the lung. The mesothelium is the single layer of cells that envelopes all lung lobes. The mesothelium, mesenchyme, and epithelium all interact through complex signaling networks to orchestrate lung development (*2, 3*).

Fibroblast Growth Factor (FGF) signaling pathways function in all stages of lung development. The FGF signaling pathway has 18 signaling ligands that activate four transmembrane tyrosine kinase receptors (FGFRs). *Fgfrs* are expressed in both mesenchyme and epithelium, and these cell-types preferentially produce alternative splice variants of FGFRs which modifies their ligand binding specificity and thus the responsiveness of these cell types to different FGF ligands (*4*). During the early pseudoglandular stage (E9.5-E14.5), FGF10 is produced in mesenchyme and FGF9 is produced in mesothelium and epithelium (*3, 5*).

FGF10 primarily signals to the b splice variant of FGFR2 (FGFR2b) in lung epithelium (*6, 7*). In the absence of FGF10 or FGFR2b, the initial lung buds form but fail to grow or branch (*8–12*). However, when FGF10-FGFR2b signaling is reduced or conditionally disrupted after initial bud formation, branching is impaired and, in some studies, epithelial cell death is increased (*13–18*). Downstream of FGF10-FGFR2b, the ERK-MAPK signaling pathway directs epithelial proliferation and bud outgrowth (*19, 20*).

*Fgf9* is expressed in lung epithelium during the first half of the pseudoglandular stage of development, from E9.5-E12.5, and in the mesothelium throughout most of the pseudoglandular stage, E9.5-E14.5 (*21–24*). Mice lacking FGF9 (*Fgf9*^−/−^) die at birth and their lungs are small with reduced mesenchyme and airway branching (*25, 26*). Investigation of the differential roles of epithelial- and mesenchymal-derived FGF9 signaling showed that mesothelial-derived FGF9 mainly functions to direct mesenchymal proliferation. Mechanistic studies identified FGFR1 and FGFR2 as the mesenchymal receptors that respond to FGF9, pErk as a downstream signal, and mesenchymal Wnt-β-catenin as a feedforward signal that is essential to maintain mesenchymal *Fgfr1* and *Fgfr2* expression (*3, 24, 27, 28*).

By contrast, inactivation of epithelial FGF9 primarily affects epithelial budding (*24*) and overexpression of FGF9 leads to epithelial dilation (*26, 29, 30*). Initial models showed that FGF9 overexpression in pseudoglandular stage epithelium induces *Fgf7* and *Fgf10* in adjacent mesenchyme, and it was hypothesized that this increase in FGF7 and FGF10 causes the observed epithelial dilation (*26, 30*). However, evidence from several studies indicated that effects of FGF9 on pseudoglandular stage epithelium may result from FGF9 signaling directly to epithelium. For example, isolated lung epithelium from E12.5 embryos directly responds to FGF9 (*30*), and lung explants lacking mesenchymal FGFR1 and FGFR2 fail to show a mesenchymal response to FGF9 (growth), but still show epithelial dilation in response to FGF9, consistent with FGF9 signaling directly to epithelium (*24, 27*). *del Moral et al* reported epithelial explants lacking FGFR2 fail to respond to FGF9, and concluded that FGF9 induces distal epithelial proliferation directly through activation of epithelial FGFR2b (*30*). However, this is inconsistent with receptor specificity studies (*6*), and with in vivo overexpression of FGF9 in lung epithelium, which results in mesenchymal hyperplasia and epithelial dilation, without affecting epithelial proliferation (*29*).

The epithelial FGFR that responds to FGF9 during pseudoglandular stage lung development is thus not known. In adult lung, overexpression of FGF9 in lung epithelium results in adenocarcinoma, without affecting lung stroma (mesenchyme) and the induction of adenocarcinoma by FGF9 was exclusively dependent on FGFR3 (*31, 32*). Collectively, these data show that FGF9 signals through FGFR1 and FGFR2 in pseudoglandular stage mesenchyme and FGFR3 in adult epithelium. Consistent with this, in vitro receptor specificity studies show that FGF9 can efficiently activate mesenchymal FGFR1c and FGFR2c and epithelial FGFR3b splice variants (*6, 33, 34*). We therefore hypothesized that FGF9 signaling to FGFR3 may also function during embryonic lung development.

Previous studies convincingly show that the original notions of strict and independent epithelial to mesenchymal and mesenchymal to epithelial FGF signaling do not account for all of the FGF signals in pseudoglandular stage lung development (*24, 27, 30*). However, why several non-redundant FGFs are required for lung epithelial development and how signaling by these different FGFs is coordinated in lung epithelial cells, is not known. Here, we examined how FGF9 and FGF10 signaling impart unique functions during pseudoglandular stage lung development. We identified a partially antagonistic relationship between FGF10-FGFR2b and FGF9-FGFR3 signaling in which FGF9 directly promotes distal epithelial fate specification and inhibits epithelial differentiation through FGFR3, whereas FGF10 promotes epithelial proliferation and differentiation through FGFR2. We further showed that these two ligand-receptor pairs rely on different downstream signaling pathways.

## Results

### FGF9 signals to lung epithelium in the absence of functional mesenchymal FGF receptors

To determine whether FGF9 can directly signal to the lung epithelium in whole-lung explants, we tested explants with genetically impaired mesenchymal FGFR1 and FGFR2 signaling for their responsiveness to FGF9. We used the *Dermo1*^*Cre*^ (*Twist2*^*Cre*^) allele to target lung mesenchyme (*26, 35*) and mated it with mice carrying floxed alleles of *β-Catenin (Ctnnb1)* or *Fgfr1* and *Fgfr2* (*35–37*). Lungs with the genotype *Dermo1*^*Cre/+*^; *β-Catenin*^*f/f*^ (Dermo1-βCat-KO), *Dermo1*^*Cre/+*^; *Fgfr1*^*f/f*^; *Fgfr2*^*f/f*^ (Dermo1-FGFR1/2-KO), and *Fgf9*^*LacZ/LacZ*^ (*Fgf9*^−/−^) (*38*) were placed in explant culture on embryonic day 11.5 (E11.5) for 48 h. Due to loss of a mesenchymal FGFR-Wnt-β-Catenin feed-forward signaling loop, all of these genotypes lack mesenchymal FGFR1 and FGFR2 signaling (*24, 27*). Wild-type explants treated with FGF9 showed mesenchymal expansion, epithelial dilation, and reduced numbers of buds (Fig. 1, A and B), consistent with previous observations (*24, 26, 27, 30*). By contrast, Dermo1-βCat-KO, Dermo1-FGFR1/2-KO, and *Fgf9*^−/−^ explants all showed an absence of mesenchymal growth in response to FGF9 but still showed marked epithelial dilation and reduced epithelial branching (Fig. 1, A, C to E). These data showed that lung epithelium could respond to FGF9 independently of mesenchymal FGF signaling in whole-organ explants.

**Fig. 1.**
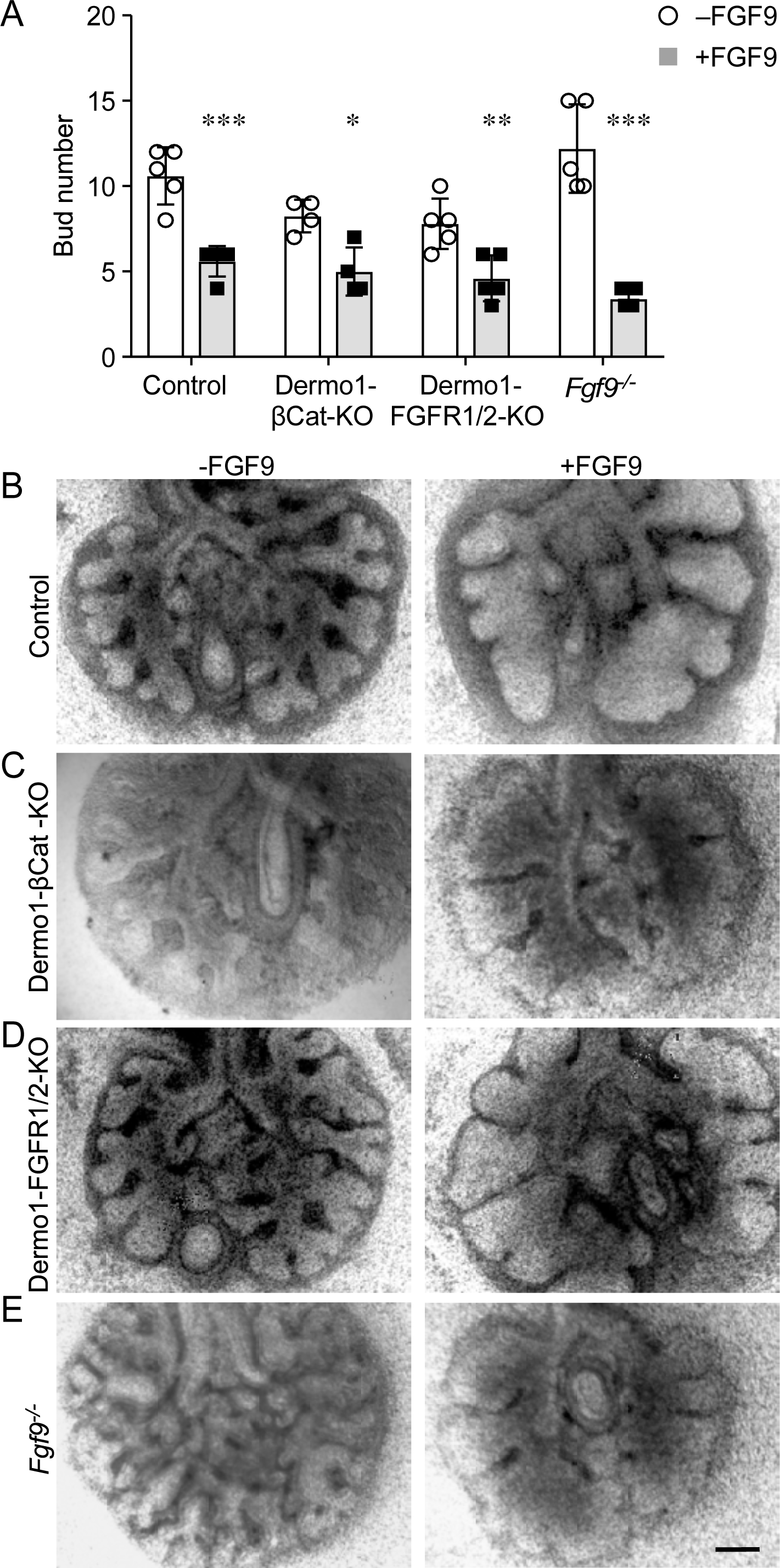
FGF9 signals to lung epithelium in the absence of functional mesenchymal FGF receptors. (**A**) Quantification of bud number in response to BSA or FGF9 in E11.5 lung explants cultured for 48 h. (**B–E**) Explant cultures from wild-type control (B), Dermo1-βCat-KO (C), Dermo1-FGFR1/2-KO (D), and *Fgf9*^−/−^ (E) mice. 2 Way ANOVA, Sidak’s multiple comparisons test, n= 4–5 explants. \**p<* 0.01, ** *p<* 0.02, *** *p<* 0.0001). Scale bar, 200 µm.

### FGF9 signals to lung epithelium through FGFR3

To further investigate whether FGF9 could directly target lung epithelium, E11.5 wild-type distal lung epithelial bud tips were isolated and cultured in Matrigel in the presence or absence of FGF9. After 48 h of culture, isolated epithelial explants failed to expand in the absence of FGF9 and formed an enlarged cyst-like structure in the presence of FGF9 (Fig. 2, A to C). Consistent with previous studies (*30*), these data show that functional FGF receptor(s) that can respond to FGF9 must be present on the surface of distal lung epithelium. Based on previous studies showing that FGF9 signals through FGFR3 in adult lung epithelium to induce adenocarcinoma and specificity studies that identify the FGF9 family as uniquely able to activate the b splice variant of FGFR3 (*6, 31–34*), we hypothesized that FGFR3 would mediate the response of embryonic lung epithelium to FGF9. Immunostaining of E12.5 and E13.5 lung sections showed the presence of both FGFR2 and FGFR3 on the basal surface of the distal developing airway epithelium and in mesenchyme (fig. S1). The similar epithelial expression pattern suggested that epithelial cells express both receptors. At E13.5, FGFR2 was also present throughout mesenchyme, while FGFR3 was expressed in a subset of lung mesenchymal cells. These data are consistent with a requirement for FGFR3 for the FGF9 response in epithelial explant cultures (Fig. 2, C and D).

**Fig. 2.**
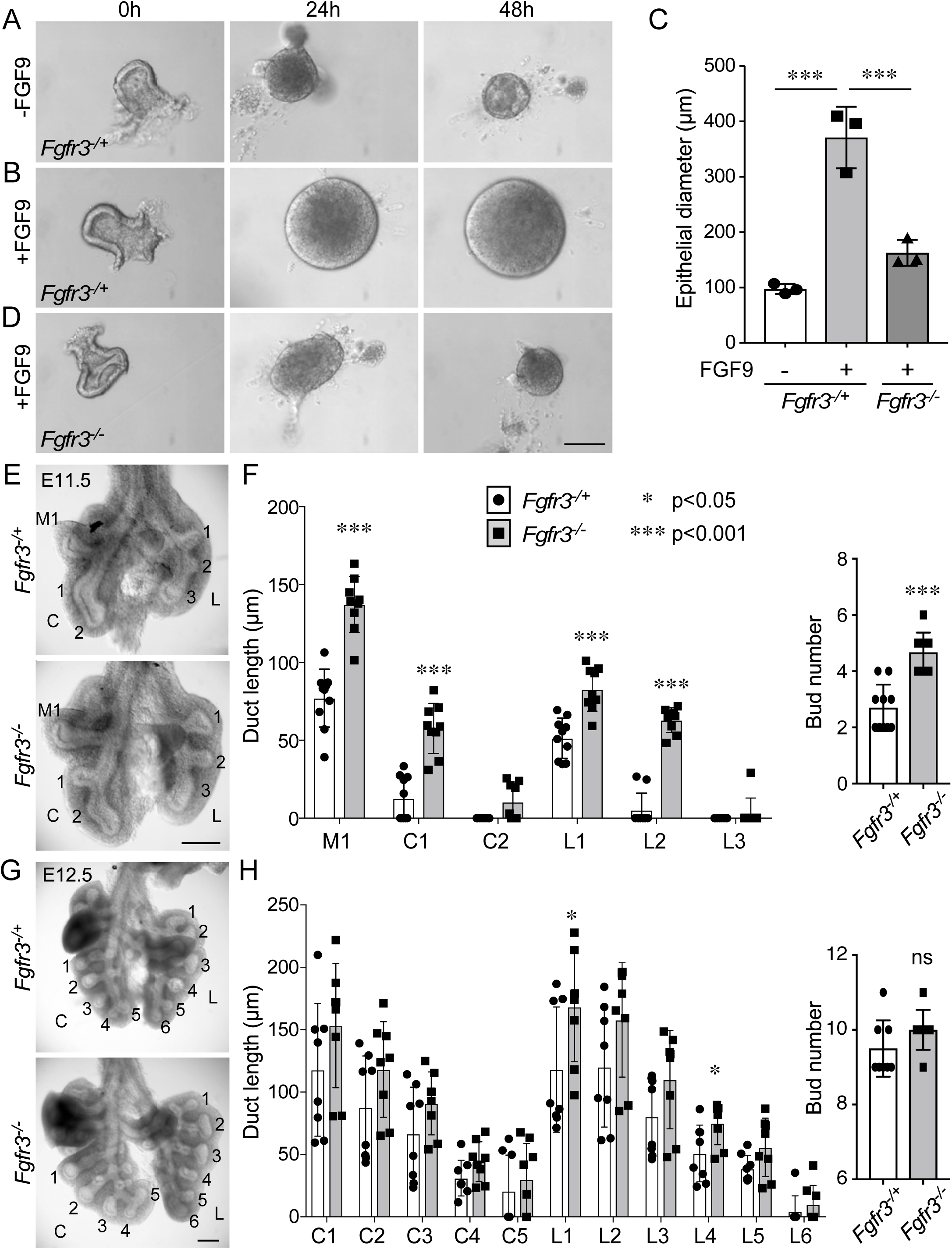
FGF9 signals to lung epithelium through FGFR3. (**A, B, D**) Isolated E11.5 distal lung epithelium from *Fgfr3*^*-/+*^ (A, B) and *Fgfr3*^−/−^ (D) embryos cultured in Matrigel in the presence or absence of FGF9 for 0, 24, and 48 hours. (**C**) Quantification of lung epithelial diameter at 48 h. ANOVA with Tukey’s multiple comparison test. n = 3 epithelial explants per genotype. (**E**) Anterior views of gross dissections of *Fgfr3*^*-/+*^ and *Fgfr3*^−/−^ lungs at E11.5. (**F**) Quantification of duct length and bud number at E11.5. All middle (M), caudal (C), and left (L) lobe duct lengths and bud numbers were counted. (**G**) Anterior views of gross dissections of *Fgfr3*^*-/+*^ and *Fgfr3*^−/−^ lungs at E12.5. (**H**) Quantification of duct length and bud number at E12.5. The caudal lobe and left lobe duct lengths and bud numbers were counted. Bud length was analyzed using the Wilcoxon rank sum test, and bud number was analyzed using the Student’s t-test. n = 9-10 lungs at E11.5 and 8 lungs at E12.5. * *p<* 0.05; *** *p<* 0.001; ns, not significant. Scale bars, 100 µm (A–C), 200 µm (E, G).

To establish the in vivo requirement for FGFR3 during early pseudoglandular stage development, lungs were isolated from E11.5 and E12.5 heterozygous (*Fgfr3*^*-/+*^) and knockout (*Fgfr3*^−/−^) embryos. Lungs from *Fgfr3*^−/−^ embryos appeared larger than those from *Fgfr3*^*-/+*^ embryos at both time points. At E11.5, quantification of bud number and length revealed increased numbers of buds and increased bud length in *Fgfr3*^−/−^ compared to *Fgfr3*^*-/+*^ lungs (Fig. 2, E and F). At E12.5, there was a trend towards increased bud length that was only significant in the left lobe (L1 and L4) ducts (Fig. 2, G and H). Bud number was not significantly different at E12.5. Consistent with this phenotype, *Fgfr3*^*-/+*^ E11.5 explants (cultured for 48 h) showed a greater decrease in bud number in response to treatment with FGF9 compared to *Fgfr3*^−/−^ explants treated with FGF9 (fig. S2, A to C). We interpret this as failure of FGF9 to suppress branching in the absence of FGFR3. In the absence of added FGF9, bud number was similar between *Fgfr3*^*-/+*^ and *Fgfr3*^−/−^ explants. These data show that FGFR3 affects lung epithelial development during early embryonic stages but that this phenotype is masked or compensated at later stages.

### FGF9 directs distal epithelial specification and differentiation but not proliferation or viability

We previously showed that lung epithelial proliferation was not significantly affected in *Fgf9*^−/−^ embryos at E10.5-E13.5 (*25*) or in FGF9-overexpressing embryos induced from E10.5-E12.5 (*29*). To confirm that FGF9 signaling does not affect pseudoglandular stage lung epithelial proliferation and to extend the analysis to include any effects on cell death, we examined the presence of phosphorylated histone H3 (PHH3) and terminal deoxynucleotidyl transferase dUTP nick end labeling (TUNEL) in control (*Fgf9*^*-/+*^) and *Fgf9*^−/−^ (*Fgf9*^*LacZ*^ allele) embryos at E10.5, E11.5, and E12.5 (fig. S3, A and B). Quantification of labeled nuclei per total nuclei in epithelium showed no significant change in cell death (Table 1A) or proliferation (Table 1B) in control and *Fgf9*^−/−^ lung at these time points.

**Table 1.** Cell death and cell proliferation in lung epithelium.

| A | TUNEL+ cells/1000 epithelial cells |  |  |
| --- | --- | --- | --- |
|  | E10.5 | E11.5 | E12.5 |
| Control | 10.6<sup>#</sup> ±6.7 | 0.0 ±0 | 0.6 ±0.5 |
| *Fgf9*<sup>-/-</sup> | 1.7 ±2.9 | 0.8 ±1.4 | 0.9 ±0.7 |
| p | 0.1 | 0.4 | 0.7 |

| B | PHH3+ cells/1000 epithelial cells |  |  |
| --- | --- | --- | --- |
|  | E10.5 | E11.5 | E12.5 |
| control | 65.8 ±29.0 | 44.5 ±32.7 | 52.6 ±23.2 |
| *Fgf9*<sup>-/-</sup> | 81.1 ±26.0 | 73.6 ±8.9 | 31.2 ±12.0 |
| p | 0.5 | 0.2 | 0.2 |

<sup>#</sup> Mean ± SD. Three control and three *Fgf9*<sup>-/-</sup> lungs were sectioned and
all epithelial cells in all sections from each lung were counted (294-1411 epithelial cells/lung).

As a result of loss of *Fgf9* not affecting epithelial proliferation or cell death, we next examined epithelial proximal-distal specification and differentiation. SOX9 marks embryonic lung epithelial progenitors that undergo branching morphogenesis. SOX9 progenitors give rise to SOX2 producing cells that will become the future conducting and alveolar airway epithelium (*39, 40*). SOX2 production therefore marks non-branching proximal lung epithelium while SOX9 production is restricted to distal branching epithelium (*41, 42*).

At E13.5, the SOX2 domain was expanded in proximal regions while the SOX9 domain was restricted to more distal regions in *Fgf9*^−/−^ lungs compared to heterozygous control lungs (Fig. 3, A and B). To test the effects of FGF9 overexpression on proximal-distal specification, Spc-rtTA; Tre-Fgf9-Ires-Gfp (SPC-FGF9-OE) mice and Spc-rtTA control mice were induced with doxycycline from E10.5-E14.5. By contrast to the loss of function experiment, when FGF9 was overexpressed, all airway epithelium produced SOX9, while SOX2 was absent from all but the very proximal tracheal epithelium (Fig. 3C). However, overexpression of FGF9 did not affect epithelial identity, as Nkx2-1 and Krt5 were distributed in all epithelial cells (fig. S4, A and B), and Trp63 positive basal cells were present in proximal airway epithelium (fig. S4C, arrows). Together, these data suggest that FGF9 may suppress the conversion of SOX9 producing progenitors to non-branching SOX2 producing respiratory epithelium during pseudoglandular stages of lung development, without affecting epithelial identity.

**Fig. 3.**
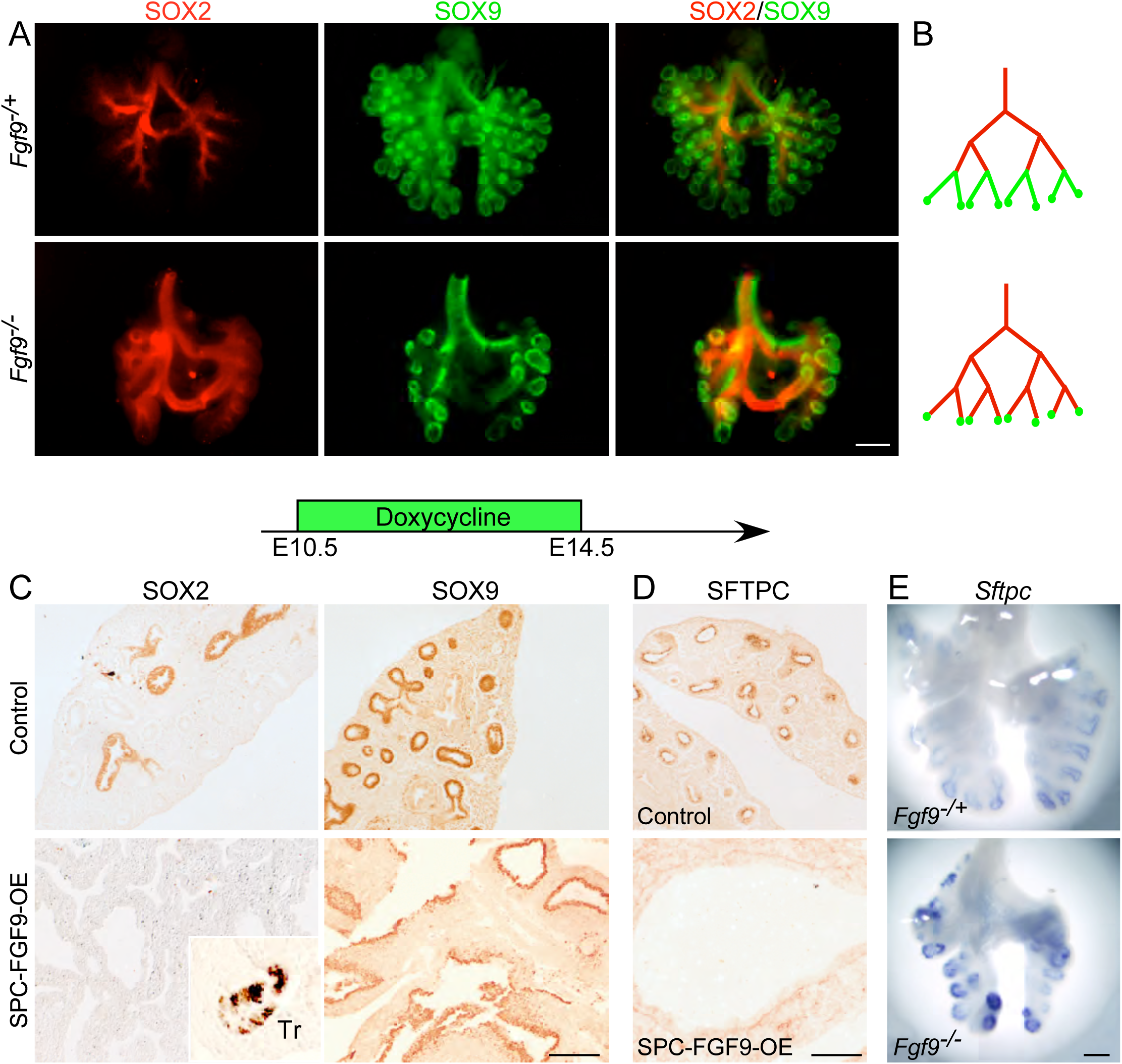
FGF9 directs distal epithelial specification and differentiation in pseudoglandular stage lung. (**A**) SOX2 and SOX9 distribution (immunofluorescence) in whole E13.5 *Fgf9*^*-/+*^ and *Fgf9*^−/−^ lungs. (**B**) Diagram of the distribution patterns shown in A. (**C, D**) SOX2 and SOX9 (C) and SFTPC (D) (immunostaining) in control (Sftpc-rtTA) and SPC-FGF9-OE lung sections that were induced with doxycycline from E10.5-14.5 and harvested at E14.5. (**E**) Comparison of *Sftpc* expression (in situ hybridization) in *Fgf9*^*-/+*^ and *Fgf9*^−/−^ lungs at E12.5. All images are representative of at least 3 embryos. Scale bars, 200 µm (A and E) and 100 µm (C and D).

Because FGFR3 is the likely receptor for FGF9, we also examined the effects of loss of FGFR3 on proximal-distal epithelial specification. Similar to the *Fgf9*^−/−^ lungs, *Fgfr3*^−/−^ lungs showed a distal expansion of the SOX2 distribution pattern (fig. S5A). However, unlike the *Fgf9*^−/−^ lungs, the SOX9 distribution pattern in *Fgfr3*^−/−^ lungs remained similar to control, resulting in an increased region of epithelium that contained both SOX2 and SOX9 (fig. S5B arrow). These data support a model in which FGF9 signals to FGFR3, but also suggests that FGF9 may transmit additional signals, either through another epithelial FGFR or, indirectly, through mesenchymal FGFR1 and FGFR2, which should still be active in the *Fgfr3*^−/−^ lungs.

We next examined the effects of FGF9 on epithelial differentiation by examining expression of *Sftpc*, an early marker of epithelial differentiation. Consistent with FGF9 functioning to suppress epithelial differentiation, induced overexpression of FGF9 from E10.5 to E14.5 inhibited all SFTPC production (Fig. 3D). By contrast, at E12.5 (Fig. 3E) distal epithelium showed increased expression of *Sftpc* in *Fgf9*^−/−^ lungs compared to *Fgf9*^*-/+*^ control lungs. Similarly, distal epithelium showed increased expression of *Sftpc* in *Fgfr3*^−/−^ lungs compared to *Fgfr3*^*-/+*^ control lungs (fig.S5C).

To determine whether FGF9 could reverse specification of distal (SOX2 producing) bronchial epithelium, SPC-FGF9-OE and Spc-rtTA control mice were induced with doxycycline from E16.5-E18.5. In control mice SOX2 was distributed in all conducting airways and SOX9 was located in very distal epithelial cells, whereas in SPC-FGF9-OE mice SOX2 was only present in the most proximal airway epithelium and the SOX9 distribution pattern was expanded to most airway epithelium (Fig. 4, A to C). Similar to earlier time points, both control and SPC-FGF9-OE lungs showed similar distribution patterns of Nkx2-1, Krt5, and Trp63 (Fig. 4, D to F). These data suggest that FGF9 can reverse bronchial epithelial specification at a late embryonic time point.

**Fig. 4.**
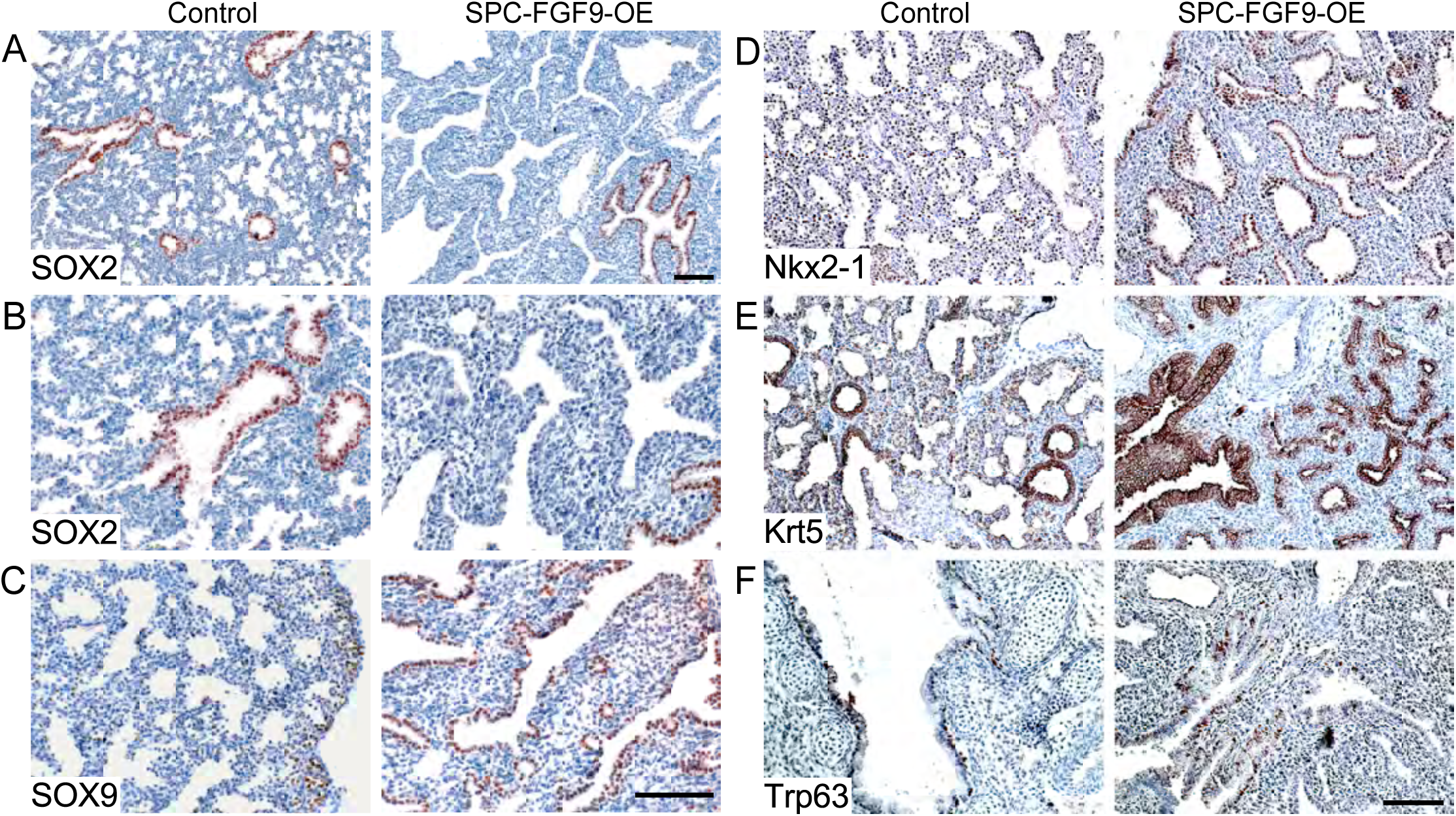
FGF9 can reverse epithelial specification without affecting epithelial identity. SPC-FGF9-OE mice and Sftpc-rtTA control mice were induced with Dox from E16.5-18.5 and lung sections were immunostained for (A, B) Sox2, (C) Sox9, (D) Nkx2-1, (E) Krt5, and (F) Trp63. All images are representative of at least 3 embryos. Scale bars, 100µm.

### FGF9 and FGF10 have distinct effects on pseudoglandular stage lung epithelium

To directly compare effects of overexpressing FGF9 and FGF10 on pseudoglandular stage lung development, we induced Spc-rtTA; Tre-Fgf10 (SPC-FGF10-OE) and SPC-FGF9-OE mice with doxycycline from E10.5 to E12.5. In both models, the overall size of the lungs was larger compared to Spc-rtTA controls (Fig. 5A). Both FGF9- and FGF10-induced lungs showed mesenchyme expansion and distal airway dilation although this was more prominent in the FGF9 overexpressing lungs. However, when proximal-distal specification was examined by immunostaining for SOX2 and SOX9, SPC-FGF9-OE lungs clearly showed expansion of SOX9-producing epithelium and loss of SOX2 producing epithelium (Fig. 3C and 5B), while SPC-FGF10-OE lungs showed relatively normal patterns of SOX2 and SOX9 producing epithelium, which were comparable to control lungs (Fig. 5B).

**Fig. 5.**
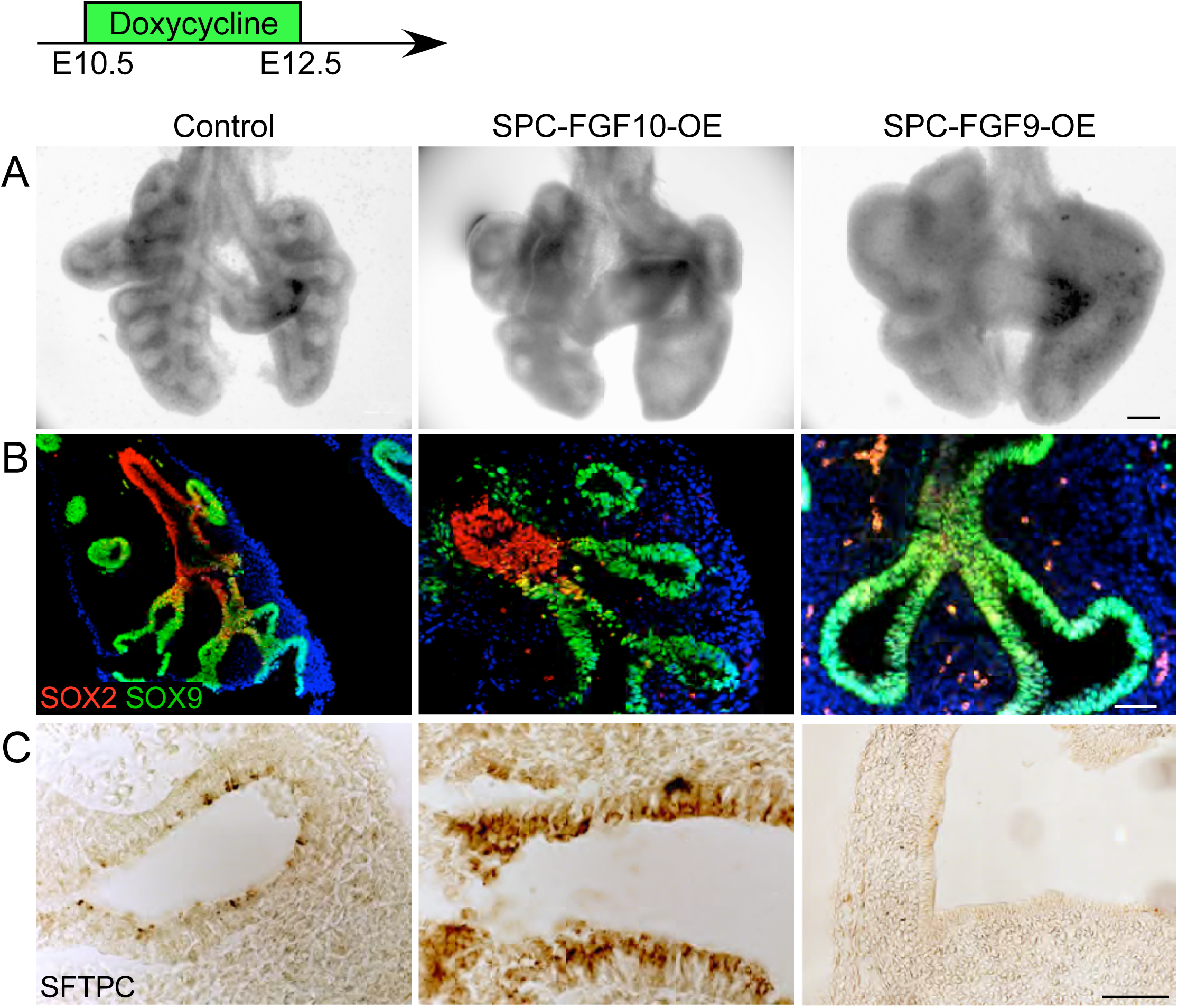
FGF9 and FGF10 have distinct effects on pseudoglandular stage lung epithelium. (**A**) Anterior views of whole E12.5 lungs. Control (Sfptc-rtTA or Tre-Fgf10 or Tre-FGf9-Ires-Gfp). All embryos were induced with doxycycline from E10.5-12.5. Note that both FGF9 and FGF10 overexpressing lungs showed mesenchyme expansion and distal airway dilation. (**B**) SOX2 and SOX9 distribution (immunofluorescence). (**C**) SFPTC immunostaining. All images are representative of at least 3 embryos. Scale bars, 200 µm (A), 100 µm (B), 50 µm (C).

We next examined the effects of FGF9 and FGF10 overexpression on epithelial differentiation by immunostaining for SFTPC. Control lungs show detectable SFTPC production at E12.5 (Fig. 5C). SPC-FGF9-OE lungs, induced with doxycycline from E10.5 to E12.5 showed absence of SFTPC production. However, SPC-FGF10-OE lungs showed normal or possibly increased SFTPC production (Fig. 5C). To complement the FGF10 activation studies, we also looked at lungs in which the FGF10 receptor, FGFR2 was inactivated in lung epithelium. Mice containing the *Shh*^*CreER*^; *ROSA*^*tdTomato*^ *(Ai9)* alleles were injected with tamoxifen at E10.5 and embryos were harvested at E13.5 (Fig. 6A). tdTomato producing cells were primarily located in distal epithelium and in epithelial tips (Fig. 6B). We then generated a *Shh*^*CreER*^; *Fgfr2*^*f/f*^ (SHH-FGFR2-CKO) mouse line in which the FGF10 receptor, FGFR2, could be conditionally inactivated in lung epithelium after the initiation of branching morphogenesis. Female mice carrying SHH-FGFR2-CKO embryos were injected with tamoxifen at E10.5 and the lungs were collected at E13.5. Compared to *Shh*^*CreER*^ control lungs, SHH-FGFR2-CKO lungs were larger, with more mesenchyme and elongated ducts (Fig. 6C). Consistent with the FGF10 overexpression data, when FGFR2 was inactivated there were minimal effects on the distribution pattern of SOX2 (Fig. 6D). The extent of SOX9 producing epithelium, however, was somewhat expanded (Fig. 6E). Epithelial differentiation, assessed by immunostaining, showed notably reduced distribution of SFTPC in SHH-FGFR2-CKO compared to control lungs (Fig. 6F).

**Fig. 6.**
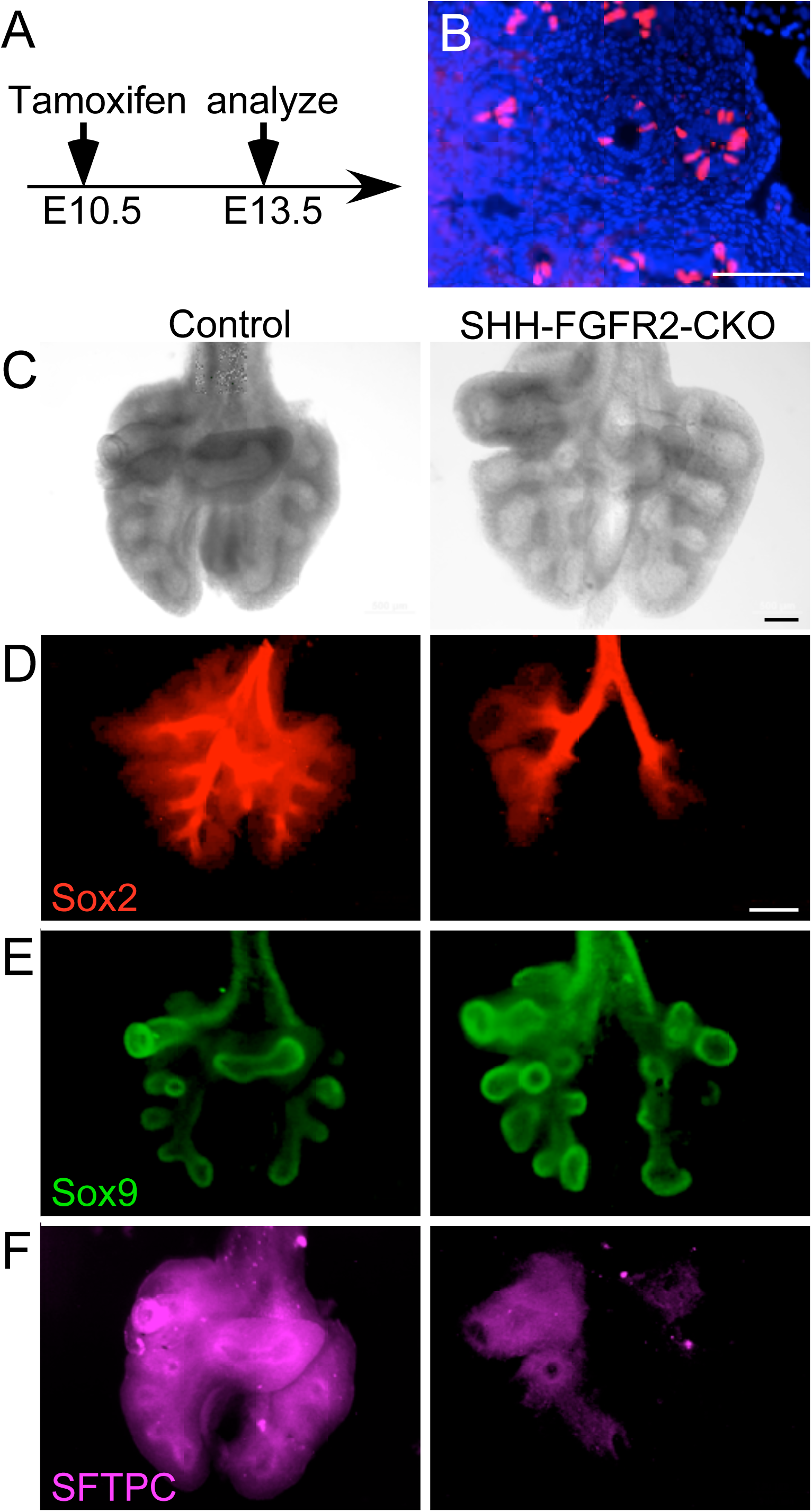
Conditional epithelial inactivation of *Fgfr2* shows similar phenotypes to *Fgf9* overexpression. (**A**) Experimental plan. All embryos were induced with Tamoxifen (i.p. injection) at E10.5 and harvested at E13.5. (**B**) Frozen section from *Shh*^*CreER*^; *ROSA*^*tdTomato*^ showing lineage labeled cells in the epithelial tips. (**C–F**) Anterior views of E13.5 whole control (*Shh*^*CreER*^ or *Shh*^*CreER*^; *Fgfr2*^*f/+*^) and lungs lacking epithelial FGFR2 (SHH-FGFR2-CKO). (C) brightfield image. (**D**) SOX2, (E) SOX9, (F) SFTPC distribution (whole mount immunofluorescence). All images are representative of at least 3 embryos. Scale bars, 100 µm (B), 50 µm (C), 200 µm (D–F).

These data indicate that FGF9 has much greater effects on epithelial specification than FGF10, with FGF9 functioning to maintain a SOX9 producing distal epithelium while FGF10 directs duct outgrowth and branching. Furthermore, the epithelial differentiation phenotype associated with inactivation of epithelial FGFR2 (reduced production of SFTPC) resembles that of FGF9 activation (reduced production of SFTPC). Together, these data suggest that FGF9-FGFR3 signaling primarily directs specification while FGF10-FGFR2b signaling primarily directs proliferation, but not specification, in pseudoglandular stage epithelium.

### FGF10-FGFR2b and FGF9-FGFR3 activate different signaling pathways

FGFRs activate four primary downstream signaling pathways: RAS/MAPK, PI3K, PLCγ and STAT, to mediate cell proliferation, migration, differentiation, and survival (*4, 43, 44*). However, most signaling studies have focused on FGFR1 and to some extent, FGFR2 and very few studies have addressed differences in downstream signaling between the four FGFRs (*43*). Because activation of FGFR2 and FGFR3 in embryonic lung epithelium has distinct outcomes, we hypothesized that these FGFRs may preferentially use different downstream signaling pathways. To address this question, we examined the response of lung explants to FGF9 and FGF10 in the presence of inhibitors of the PI3K, MAPK, and STAT3 signaling pathways.

Wild-type lung explants were placed in culture on embryonic day 11.5 (E11.5) for 48 h. Over this period the explants increase in size and epithelial ducts branch and extend distally (Fig. 1A and 7A). As shown in several studies and here, treatment of explants with FGF9 resulted in mesenchymal growth, reduced branching, and epithelial dilation (Fig. 1A, 7A) (*24, 26, 27, 30, 45, 46*) while treatment with FGF10 resulted in increased branching (Fig. 7A) (*8, 45–48*). In the presence of the MEK inhibitor, U0126, which blocks MAPK mediated pErk signaling, explants showed reduced branching and epithelial dilation (Fig. 7B), a phenotype that resembles that of wild-type explants treated with FGF9 (Fig. 7A). By contrast, treatment of explants with a PI3K inhibitor, Ly2294002, resulted in increased branching (Fig. 7C), a phenotype that resembles that of wild-type explants treated with FGF10 (Fig. 7A). By contrast, the MEK inhibitor, U0126, had little effect on FGF9 which still induced epithelial dilation and suppressed branching, but strongly inhibited FGF10-induced branching (Fig. 7B). The PI3K inhibitor, Ly2294002, suppressed the epithelial dilation induced by FGF9 and had very little effect on branching induced by FGF10 (Fig. 7C). Of note, FGF9 also strongly suppressed the increased branching induced by the PI3K inhibitor (Fig. 7C), suggesting more complexity in the downstream signaling pathways activated by FGF9 or indirect effects of FGF9 signaling through mesenchyme. Treating wild-type explants with both MEK and PI3K inhibitors suppressed both epithelial branching and dilation (Fig. 7D) and caused epithelial cell death (Fig. 7, E and F). STAT3 is produced in the developing lung (*20, 49*), and can be activated by FGFRs (*50*). In the presence of the STAT3 inhibitor, Stattic (*51*), control explants showed a modest increased branching, with little effect of added FGF10 (fig. S6, A and B). STAT3 inhibition also had little effect on FGF9, as FGF9 blocked branching and induced epithelial dilation both in the presence and absence of Stattic (fig. S6, A and C).

**Fig. 7.**
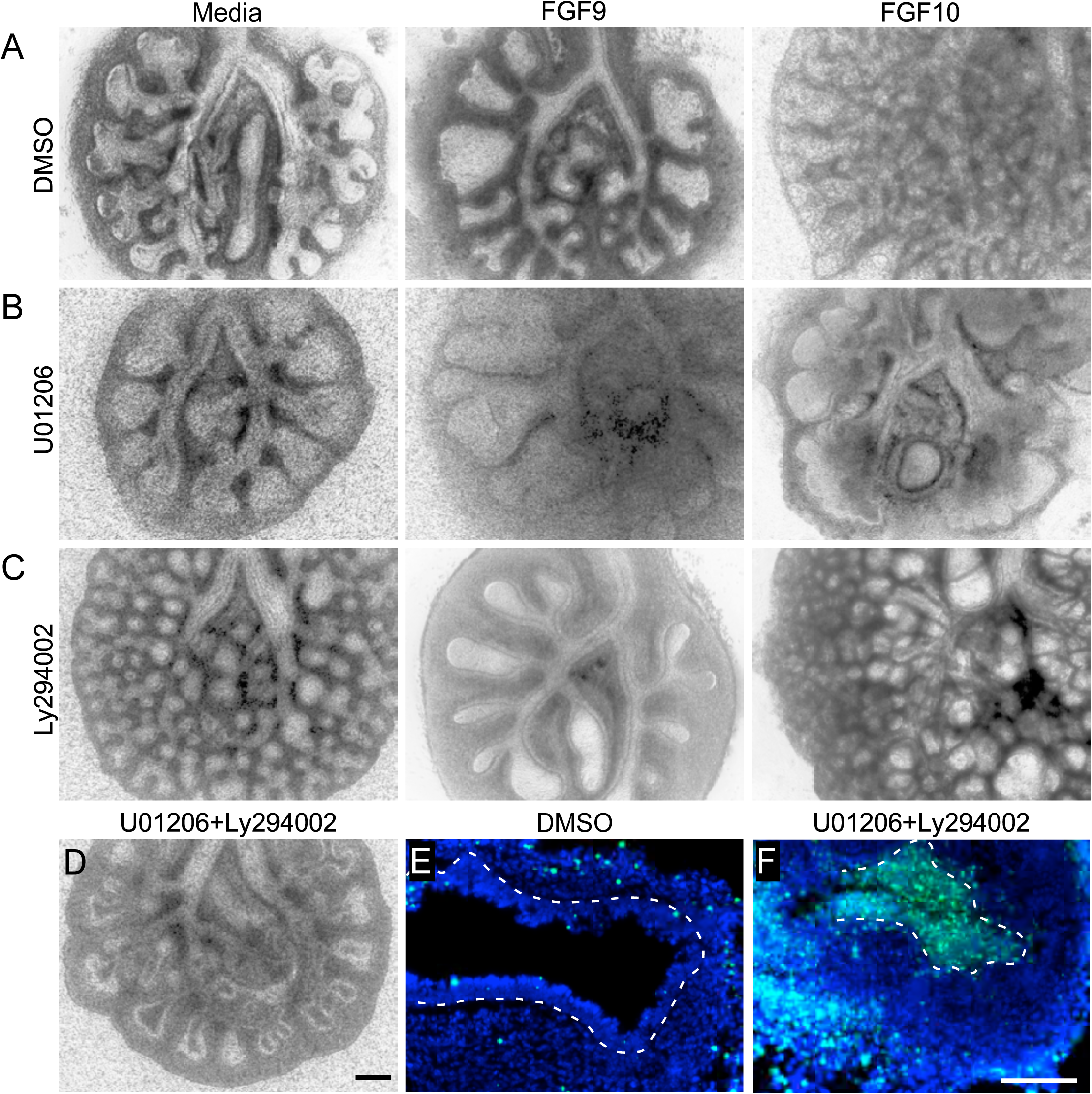
Preferential use of distinct cellular signaling pathways by FGF9 and FGF10. (A-C) E11.5 wild-type lung explants cultured for 48 h. (A, left) Controls cultured in the presence of DMSO and media lacking FGF, (A, middle) FGF9, (A, right) FGF10, (B, left) MEK inhibitor, U01206, (B, middle) FGF9 and U01206, (B, right) FGF10 and U01206, (C, left) PI3K inhibitor, Ly294002, (C, middle) FGF9 and Ly294002, (C, right) FGF10 and Ly294002. (D-F) E11.5 wild-type lung explants cultured for 48 h and treated with U01206 and Ly294002 (D and F) or DMSO (E). Brightfield image of whole lung explant (D), TUNEL stained sections (E and F). All images are representative of at least 3 embryos. Scale bars, 200 µm (A–J), 100 µm (K, L).

To further examine downstream signaling, epithelial explants from Dermo1-FGFR1/2-KO mice were treated with FGF9 or FGF10 (Fig. 8A), sectioned and then immunostained for dpERK and pAKT (Fig. 8, B and C). Treatment with FGF9 showed little effect on dpERK immunostaining but a robust induction of pAKT. By contrast, treatment with FGF10 robustly induced dpERK but had minimal effects on pAKT. These data support a model in which different FGFRs activate different downstream signaling pathways in lung epithelial cells to induce distinct developmental responses.

**Fig. 8.**
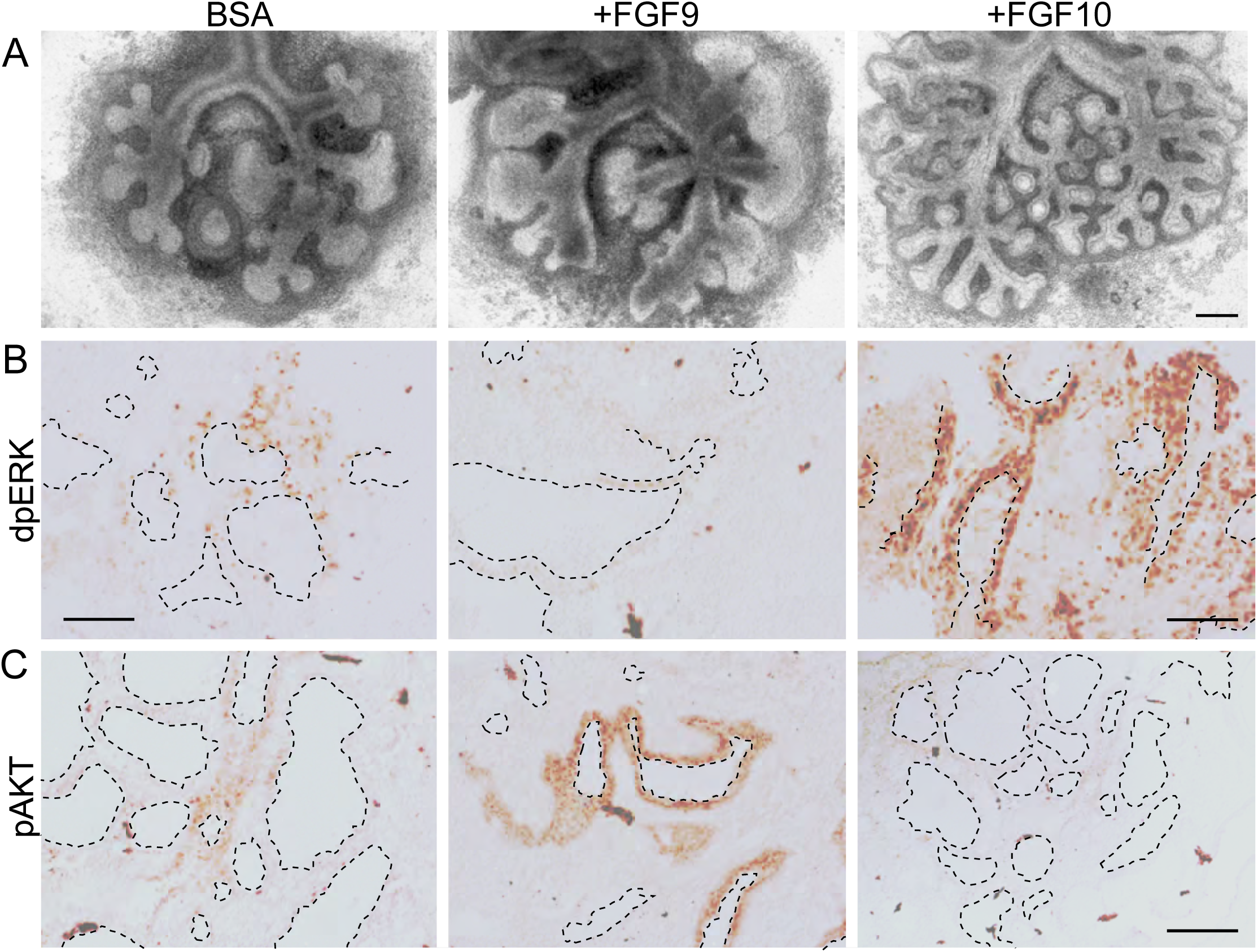
Activation of pAKT and dpERK signaling by FGF9 and FGF10 in lung explants lacking mesenchymal FGFR1 and FGFR2. (A) Images of E11.5 whole lung explants from Dermo1-FGFR1/2-KO mice cultured for 48 h and treated with BSA, FGF9, or FGF10. (B, C) Sections of lung explants immunostained for dpERK (B) and pAKT (C). All images are representative of at least 3 embryos. Dashed lines outline ductal lumens. Scale bars, 200 µm (A); 100 µm (B, C).

**Fig. 9.**
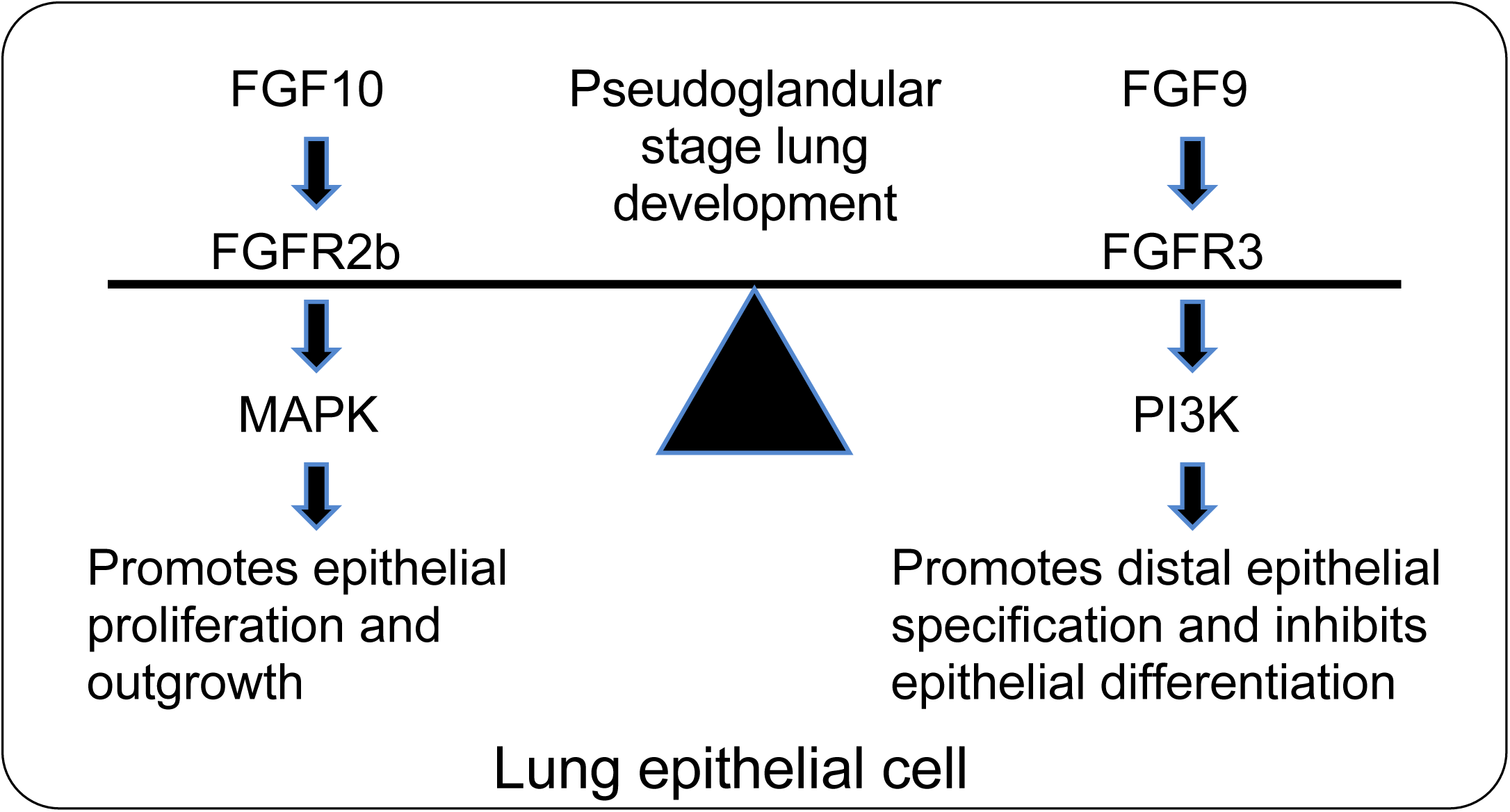
Model for FGF9 and FGF10 signaling during pseudoglandular stage lung development. FGF10 signaling through FGFR2b preferentially activates pERK and is in balance with FGF9 signaling through FGFR3 which preferentially activates PI3K.

## Discussion

Development of the lung airways involves epithelial growth, branching, and differentiation, all of which must be coordinated to make a functional lung. Many of the signals that direct lung epithelial development arise from mesenchyme; however, several studies have identified factors that act as autocrine or juxtacrine signals within lung epithelium (*3*). For example, *Bmp4* is expressed in pseudoglandular stage epithelial tips and directs epithelial proliferation and proximal-distal patterning (*52–54*). *Wnt7b* is also expressed in lung epithelium where it directs canonical Wnt signaling and expression of *Bmp4* and *Fgfr2* (*55, 56*). Here, we focused on FGF9 and FGF10, factors that are produced in developing airway epithelium and mesenchyme, respectively, that signal to epithelium to direct proximal-distal specification, differentiation, branching, and outgrowth.

### FGF9 signals through FGFR3 in early pseudoglandular stage lung epithelium

Although it has been established that FGF9 signals to pseudoglandular stage lung epithelium, the receptor that responds to FGF9 has not been rigorously defined. Initial studies suggested that FGFR2b is responsible for the FGF9 response because *Fgfr2*^−/−^ lung explants failed to respond to FGF9 (*30*). However, in vitro, FGF9 does not bind to FGFR2b, and in vivo FGFR2b is an established receptor for FGF10, which has very distinct effects on lung epithelium (proliferation and outgrowth) from that of FGF9 (specification and differentiation). The data showing that FGF9 induces epithelial proliferation in lung explants from *Fgf10*^−/−^ embryos can be explained by indirect effects through induction by FGF9 of *Fgf7* expression in adjacent mesenchyme and FGF7 activation of epithelial FGFR2b. Consistent with this model, *del Moral et al.* (*30*) showed that FGF9 strongly induces *Fgf7* expression (287%) in primary lung mesenchyme, *White et al*. (*26*) showed that overexpression of FGF9 *in vivo* in lung epithelium induces both *Fgf7* and *Fgf10* in adjacent mesenchyme, *Lebeche et al.* (*57*) showed that both *Fgf7* and *Fgf10* are expressed in primary lung bud mesenchyme and are important for growth of the lung bud, and *Bellusci et al.* and *Park et al.* showed that FGF7 is a stronger inducer of epithelial proliferation than FGF10 (*8, 58*). Additional explanations for why *Fgfr2*^−/−^ lung explants failed to respond to FGF9 include the possibility that: 1) the *Fgfr2b*^−/−^ tissue used was compromised due to lack of earlier FGF10-FGFR2b signaling; 2) FGFR3 is required for FGF9 to signal through FGFR2b by, for example, forming heterodimers; or 3) FGFR2b signaling is required for production of FGFR3 in lung epithelium.

Here, using isolated epithelium and using whole lung explants deficient in mesenchymal FGFR signaling, we confirmed that FGF9 signals directly to pseudoglandular stage lung epithelium. Using isolated epithelia that lacks FGFR3, we showed that the epithelial response to FGF9 is dependent on FGFR3, we show that *Fgfr3*^−/−^ lungs have increased duct length and bud number at E11.5, and finally, that the ability of FGF9 to suppress branching is reduced in *Fgfr3*^−/−^ explants. At E12.5, there remains a trend towards increased duct length and *Fgfr3*^−/−^ lungs appear larger than heterozygous controls. However, *Fgfr3*^−/−^ mice are viable and, as adults, do not have identified defects in lung development, although FGFR3 together with FGFR4 is important for postnatal alveologenesis (*59–61*). It is therefore likely that the *Fgfr3*^−/−^ embryonic phenotype is compensated or masked by other factors later in development. Furthermore, in the developing lung, epithelial expression of *Fgf9* is only detected at E10.5-E12.5 and is restricted to mesothelium at later times (*21–24*). This suggests that epithelial FGF9 is only needed during the early pseudoglandular stage to direct epithelial development and that mesothelial FGF9 continues to function throughout the pseudoglandular stage to direct mesenchymal development.

### FGF9 directly promotes distal epithelial fate specification and inhibits epithelial differentiation

Here, we show that inactivation of FGF9 does not affect epithelial proliferation or death during pseudoglandular stage development. We therefore hypothesized that FGF9 may direct some aspect of epithelial specification and differentiation, and that the relative distribution of SOX2 and SOX9, markers of proximal and distal lung epithelium, respectively, would be affected by the level of FGF9 signaling in epithelium. In support of this hypothesis, *Fgf9*^−/−^ and FGF9 overexpression mouse models clearly show that FGF9 directs the proximal-distal distribution of SOX2 and SOX9 expressing epithelial cells without affecting proliferation or cell death. Specifically, FGF9 overexpression expands the population of SOX9 producing epithelial progenitors, and prevents these cells from producing SFTPC, a marker of differentiation. By contrast, FGF10 overexpression or FGFR2 inactivation in epithelium had little effect on the distribution of SOX2 and SOX9 expressing epithelial cells. Although FGFR2 signaling was required to maintain SFTPC production. By contrast to our results with inducible *Fgfr2* inactivation in distal lung epithelium beginning at E10.5, *Abler et al.* (*13*) reported that inactivation of *Fgfr2* with Sftpc-Cre results in smaller lungs, increased epithelial death, reduced *Sox9* and expansion of the *Sox2* expression domain. However, unlike the limited and delayed gene targeting with *Shh*^*CreER*^ and a single tamoxifen injection at E10.5, inactivation of *Fgfr2* with Sftpc-Cre efficiently targets epithelium beginning at E9.75 (*13, 62*). It is thus likely that inactivation of *Fgfr2* at an earlier developmental time leads to epithelial apoptosis and secondary effects on proximal-distal patterning, one of which could include loss of *Fgf9* expressing cells in distal lung epithelium.

Consistent with FGF9 promoting SOX9 producing epithelium and suppressing differentiation, when *Sox9* was inactivated in epithelium using *Shh*^*Cre*^, *Sftpb* expression was increased in distal epithelium as early as E11 and there was no effect on epithelial proliferation at E13 (*63*). Of note, *del Moral et al.* (*30*) did not observe a change in SFTPC production in E12.5 explants that were cultured for 48 h in the presence of FGF9; however, because *Sftpc* expression initiates as early as E10, this explant stage may be too late to see a suppressive effect of FGF9 on *Sftpc* expression. In a more recent paper, *Jones et al.* (*18*) showed that inhibiting FGF10 signaling at E12.5 for 9 h by overexpression of a soluble FGFR2b ligand trap led to reduced expression of *Sftpc*, like what we observed with conditional inactivation of FGFR2 in epithelium from E10.5 to E13.5. Together these data support a model in which FGF9 suppresses distal epithelial differentiation.

### An antagonistic relationship between FGF10–FGFR2 and FGF9–FGFR3 signaling in the early pseudoglandular stage lung

Previous work comparing FGFR signaling in vitro suggested that differences between receptors are most likely due to differences in the strength of the tyrosine kinase domain rather than qualitative differences in downstream signaling pathways (*50, 64*). Comparison between mouse lines that affect FGF9–FGFR3 signaling and FGF10–FGFR2 signaling reveal distinctly different and, in some cases, opposite phenotypes, suggesting that these ligand-receptor pairs activate distinct signaling pathways in vivo (Fig. 8). To investigate potential differences in FGF9 and FGF10 signaling on three of the major FGFR signaling pathways, RAS–MAPK, PI3K, and STAT, we tested the response of lung explants to pathway inhibitors and treatment with FGF9 or FGF10. This analysis showed that inhibition of RAS–MAPK signaling blocks FGF10 induced branching and results in phenotypes that resemble FGF9 activation, whereas inhibition of PI3K blocks the epithelial dilation induced by FGF9 and in the absence of FGF9 results in a phenotype that resembles FGF10 activation (increased branching). However, in the presence of the PI3K inhibitor, LY294002, FGF9 still potently suppressed branching, suggesting that at the concentrations used, FGF9 effectively competes with this inhibitor or signals through additional pathways, such as the STAT3 signaling pathways. However, inhibition of STAT3, did not affect FGF9 induced epithelial dilation or suppression of branching.

The experiments with FGF9 and inhibitors were complicated by the fact that FGF9 also signals to lung mesenchyme which could activate indirect signals that affect epithelial development. However, the effects of FGF9 on epithelium were seen in both wild-type lung explants and in explants lacking mesenchymal FGFR signaling. Together, these observations suggest a model in which, in epithelium, FGFR2 and FGFR3 are activated by FGF10 and FGF9, respectively, and preferentially activate distinct downstream signaling pathways. Finally, the effects of FGF9–FGFR3 signaling and FGF10–FGFR2b signaling appear to functionally oppose each other with respect to epithelial branching, proximal-distal specification, and differentiation.

## Materials and Methods

### Mice

All mouse strains, *Fgf9*^*LacZ*^, *Fgf9*^*f/f*^, *Fgfr1*^*f/f*^, *Fgfr2*^*f/f*^, *Fgfr3*^−/−^, *Dermo1*^*Cre/+*^, *Shh*^*CreER/+*^, Spc-rtTA, Tre-Fgf9-Ires-Gfp, Tre-Fgf10, have previously been described (*26, 35–38, 65–69*). We refer to the *Fgf9*^*LacZ*^ allele as *Fgf9*^*-*^. Mice were of mixed sexes. *Fgfr3*^−/−^, Spc*-*rtTA, Tre-Fgf9-Ires-Gfp, and Tre-Fgf10 mice are on an FVB/NJ inbred background. *Fgf9*^*LacZ*^ (*Fgf9*^*-*^), *Shh*^*CreER/+*^; *Fgfr2*^*f/f*^, *Dermo1*^*Cre/+*^; *Fgfr1*^*f/f*^; *Fgfr2*^*f/f*^ mice are maintained on a mixed C57BL/6J x 129X1/SvJ genetic background. Wild-type lung explants are from mixed genetic backgrounds derived from C57BL/6J, 129X1/SvJ, and FVB/NJ.

Mice were group housed with littermates, in breeding pairs, or in a breeding harem (2 females to 1 male), with food and water provided ad libitum. Mice were housed in a specific pathogen-free facility and handled in accordance with standard use protocols, animal welfare regulations, and the NIH Guide for the Care and Use of Laboratory Animals. All studies performed were in accordance with the Institutional Animal Care and Use Committee at Washington University in St. Louis (protocol #20160113, 20190110).

### Antibodies

The following antibodies were used for immunostaining: rat anti-phospho-histone H3 (pHH3, 1:500, #H9908, Millipore-Sigma), rabbit anti-SRY-box-containing gene 9 (SOX9, 1:2000, #AB5535, Millipore Sigma), goat anti-SRY-box-containing gene 2 (SOX2, 1:250, #SC-17320, Santa Cruz Biotechnology), proSP-C (SPC, 1:500, #AB3786, Millipore-Sigma), Phospho-Akt (p-AKT, 1:150, #9271, Cell Signaling Technology), Phospho-p44/42 MAPK (dpERK, 1:200, #9101s, Cell Signaling Technology).

### Section and whole-mount immunostaining

Lung tissue was dissected and fixed following published protocols (*41, 63, 70*). Dissected embryo torsos were opened along the sternum and fixed in phosphate-buffered saline (PBS, pH 7.4) with 0.5% paraformaldehyde (PFA, #15714-S, Electron Microscopy Sciences) for 2 h for E12.5 and younger embryos and 3 h for E14.5 embryos on a rocker at room temperature. Embryos were then washed with PBS and lungs were dissected, photograph and used for cryo-section or whole-mount immunostaining. Images of whole lungs were captured on a stereo microscope (Olympus SZX12-ILLD100, Olympus).

### Lung explant cultures

Lung explant cultures were performed as described (*26*). E11.5 embryonic lungs were dissected and cultured on Transwell filters (#3462, Corning Life Sciences) for 48 h in DMEM, 2 µg/ml heparin, at 37°C, 5% CO2. Tissues were treated with mouse or human FGF9 (#100-23, #100-23, PeproTech Inc.) or human FGF10 (#100-26, PeproTech Inc.) at 2.5 ng/ml, or with inhibitors. MEK inhibitor (U01206, #9903, Cell Signaling), 25 µM; PI3K inhibitor (Ly294002, #9901 Cell Signaling), 25 µM; STAT3 inhibitor (Stattic, #S7947, Millipore-Sigma), 10 µM. Lung explants were photographed on a stereo microscope (Olympus SZX12-ILLD100) or inverted microscope (Leica, DM IL LED). Duct length was measured using ImageJ software. Data shown is representative of at least three independent experiments.

### Lung epithelial cultures

Lung epithelial cultures were performed as described (*30*). E11.5 mouse lungs were isolated and treated with undiluted dispase (#354235, Corning Life Science) for 30 min at 4°C. The distal epithelium was separated from the mesenchyme using fine tungsten needles and transferred into Matrigel™ (#356231, Corning Life Science) with culture medium (DMEM, 10% FBS, penicillin-streptomycin, L-glutamine) in a 4-well Nunc plate (#176740, ThermoFisher). Polymerization of Matrigel™ was initiated at 37°C for 30 min. After polymerization, epithelial explants were treated with either medium or medium containing recombinant FGF9 (2.5 ng/ml) and were cultured for 48 h at 37 °C in a 5% CO_2_ incubator. Explants were photographed on a stereo microscope (Olympus SZX12-ILLD100) or inverted microscope (Leica, DM IL LED). Epithelial diameter was measured using ImageJ software.

### Cell death analysis

Cyosectioned slides (7 µm), prepared as described above, were assayed by DeadEnd™ Fluorometric TUNEL System (#G3250, Promega). Slides were mounted with DAPI containing Vectashield mounting medium (#H-1200, Vector Laboratories) for fluorescent detection. Three sections, 10 µm apart, were counted for three lungs at each time point.

### In situ hybridization

The *Sftpc* (*Spc*) probe (*71*) was synthesized and labeled with a kit from Roche Applied Science. Whole mount in situ hybridization was performed as described (*26, 27*). Following color reaction and methanol dehydration lungs were photographed on an Olympus SZX12-ILLD100 stereo microscope.

## Statistical analysis

For quantification of lung tissue and explant morphologies at least 4 individual samples were measured. Data was analyzed using the unpaired Student**’** s t test, ANOVA, or Wilcoxon rank-sum test (see below). The data are reported as the mean ± SD and changes with p values less than 0.05 were considered to be statistically significant.

Duct length was compared in *Fgfr3*^−/−^ animals and *Fgfr3*^*-/+*^ lungs at each lobe/duct location using a nonparametric Wilcoxon rank-sum test. This test was chosen because the data are underlying continuous but distinctly non-Gaussian. Specifically, in most lobe/duct locations the values are skewed to the left and some have a floor of a few 0 lengths. The data include repeated measures within animals (multiple lobe and duct locations). They are not well suited to multi-level or repeated measures modeling. E11.5 data has 7 and E12.5 has 11 simultaneous Wilcoxon tests.

## Supplementary Materials

Fig. S1. FGFR2 and FGFR3 are expressed in embryonic airway epithelium.

Fig. S2. Reduced suppression of branching in response to FGF9 in explants lacking FGFR3.

Fig. S3. Loss of Fgf9 does not affect epithelial cell death or proliferation.

Fig. S4. Overexpression of Fgf9 does not affect epithelial cell identity or basal cell formation.

Fig. S5. FGFR3 directs distal epithelial specification and differentiation in pseudoglandular stage lung.

Fig. S6. Inhibition of STAT3 has minimal effects on FGF9 and FGF10 signaling.

## Acknowledgements

We thank L. Li for technical help, K. Trinkaus and the Siteman Cancer Center Biostatistics Shared Resource (https://siteman.wustl.edu/research/shared-resources-cores/biostats-core/) for statistical consultation, J. Que for providing initial embryos with epithelial *Fgfr2* inactivation. A. Hagan and K.V. Woo for critically reading the manuscript.

## Author contributions

Conceptualization, Y.Y. and D.M.O.; Methodology, Y.Y. and D.M.O.; Investigation, Y.Y. and D.M.O.; Writing Original Draft, Y.Y. and D.M.O.; Writing Review & Editing, Y.Y. and D.M.O.; Funding Acquisition, D.M.O.; Resources, D.M.O.; Supervision, D.M.O.

## Data and Materials Availability

All data needed to evaluate the conclusions in this paper are present in the paper or the Supplementary Materials.

