## Supplemental Figures S1-S6 for "FGF9 and FGF10 use distinct signaling pathways to direct lung epithelial specification and branching"

**Supplementary Materials**

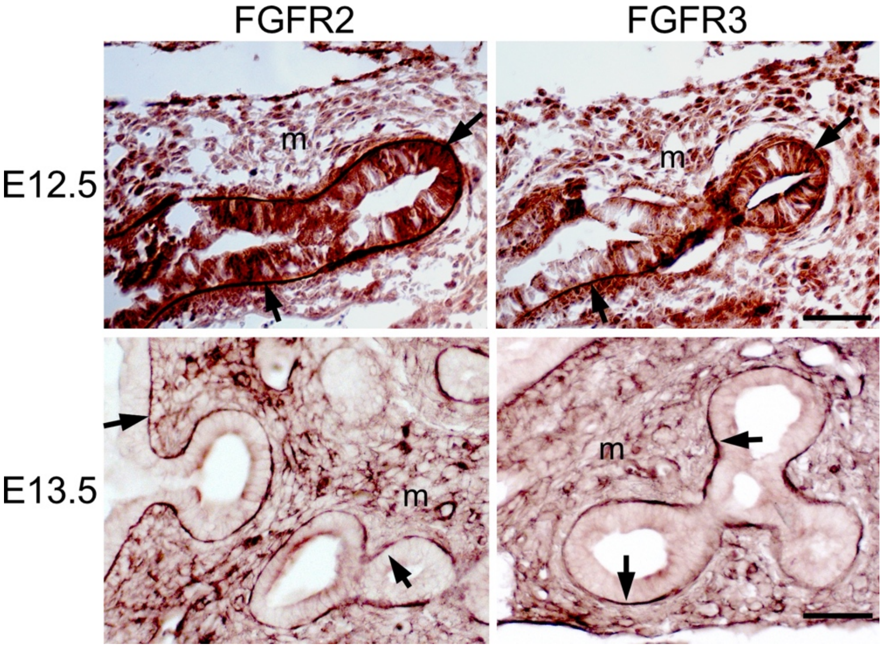


**Fig. S1. FGFR2 and FGFR3 are expressed in embryonic airway epithelium.**

Immunostaining for FGFR2 and FGFR3 on E12.5 and E13.5 lung sections. Adjacent sections are shown at E12.5. Arrows indicate staining of the basal surface of developing airway epithelium. Images representative of at least 3 embryos. m, mesenchyme. Scale bars, 50 µm.

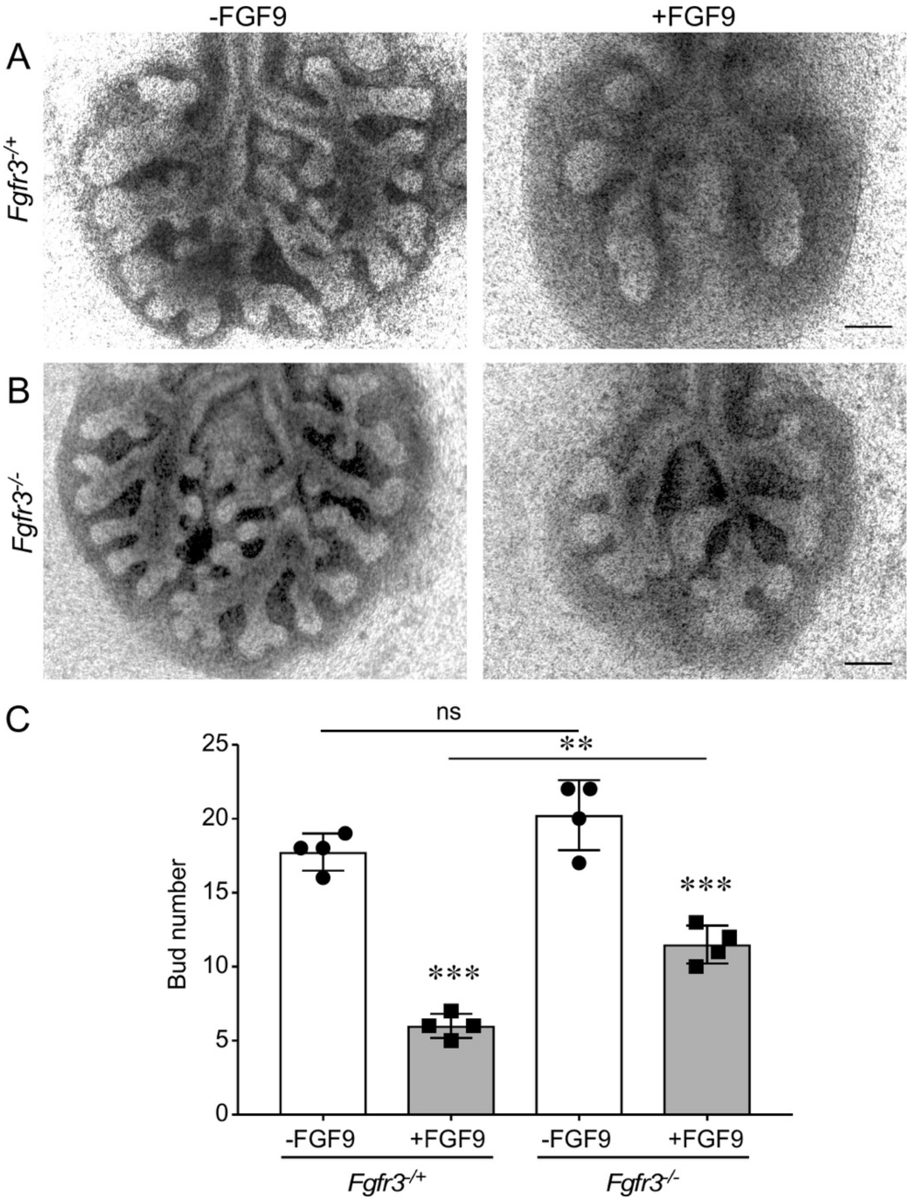


**Fig. S2. Reduced suppression of branching in response to FGF9 in explants lacking FGFR3.** (A, B) Control (*Fgfr3*$^{-/+}$) and *Fgfr3*$^{-/-}$ lung explants, cultured from E11.5-13.5 cultured in the presence or absence of FGF9. (C) Quantification of bud number in lung explants. One way ANOVA with Tukey's multiple comparisons test. n=4; ns, not significant; ** p< 0.002; *** p< 0.001. Scale bars, 200 $\mu$m.

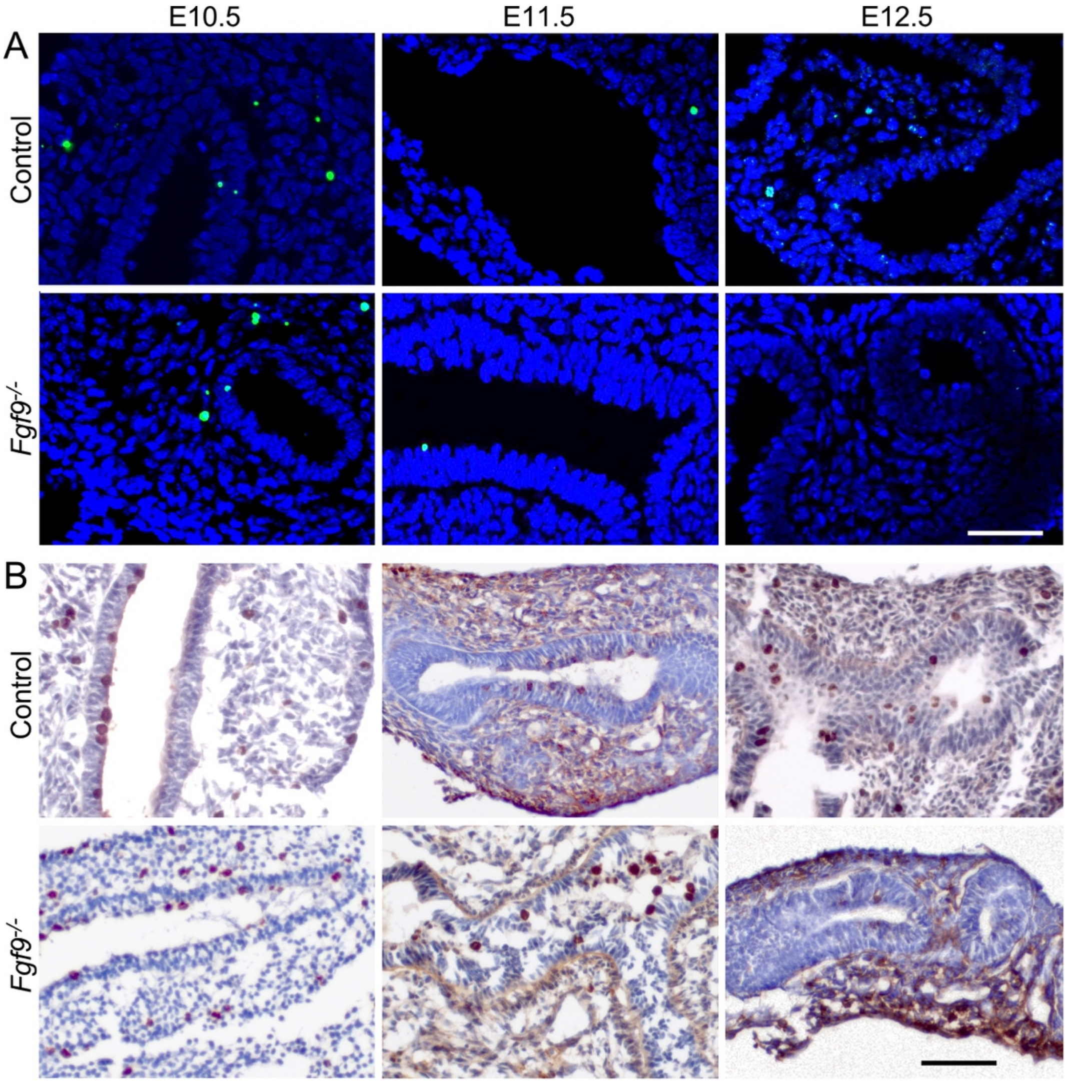


**Fig. S3. Loss of Fgf9 does not affect epithelial cell death or proliferation.** (A) TUNEL staining of histological section of control (*Fgf9*$^{-/+}$) and *Fgf9*$^{-/-}$ lungs at E10.5, E11.5 and E12.5. (B) PHH3 immunostaining of histological section of control (*Fgf9*$^{-/+}$) and *Fgf9*$^{-/-}$ lungs at E10.5, E11.5 and E12.5. Images representative of at least 3 embryos. Scale bars, 50$\mu$m.

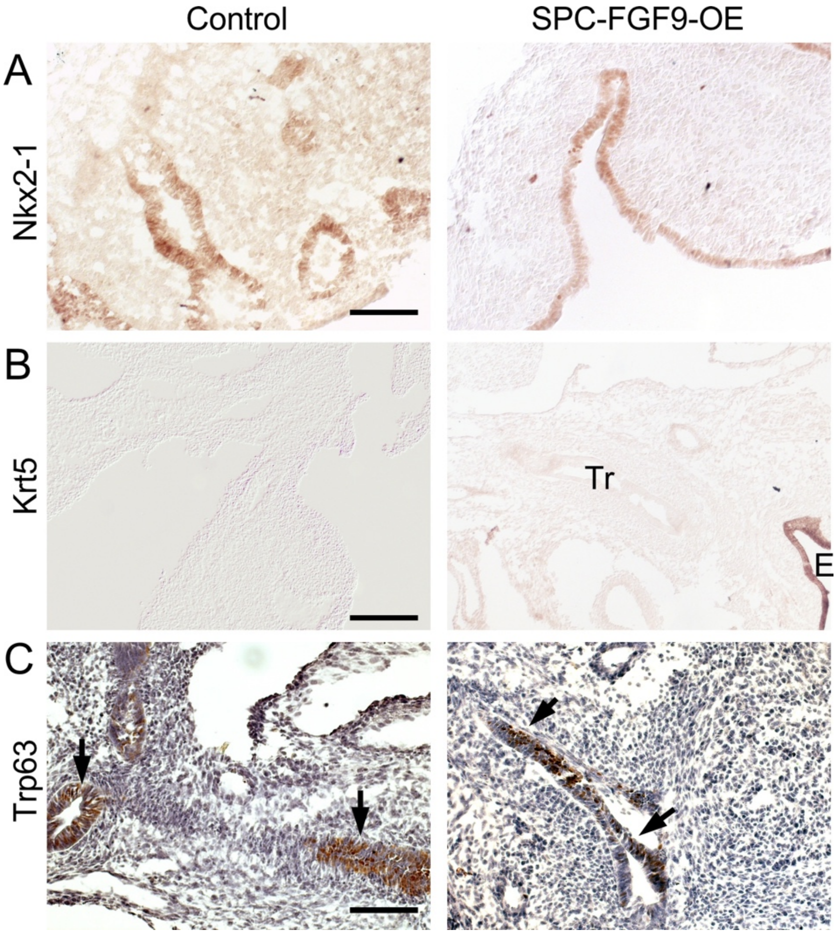


**Fig. S4. Overexpression of Fgf9 does not affect epithelial cell identity or basal cell formation.** SPC-FGF9-OE mice were induced with Dox from E10.5-12.5 and lung sections were immunostained for (A) Nkx2-1, (B) Krt5, and (C) Trp63. Arrows indicate proximal epithelium. Tr, Trachea; E, esophagus. Images representative of at least 3 embryos. Scale bar, A, C, 100μm; B, 200 μm.

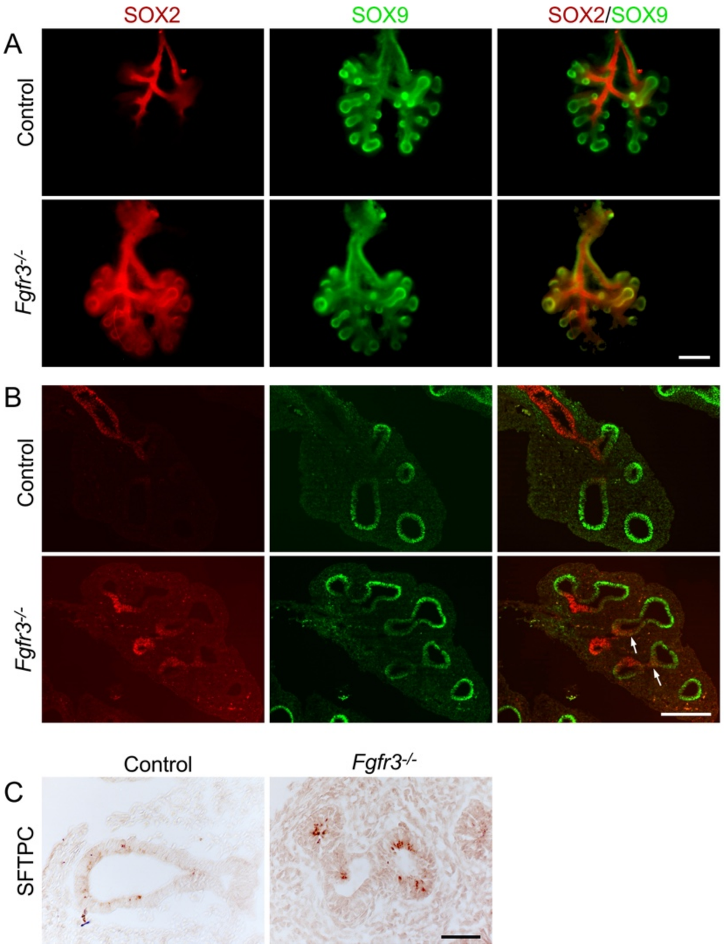


**Fig. S5. FGFR3 directs distal epithelial specification and differentiation in pseudoglandular stage lung. (A, B)** SOX2 and SOX9 distribution (immunofluorescence) in (A) whole and (B) histological sections of E12.5 control (*Fgfr3*$^{-/+}$) and *Fgfr3*$^{-/-}$ lungs. (C) SFTPC immunostaining of histological sections of E12.5 control (*Fgfr3*$^{-/+}$) and *Fgfr3*$^{-/-}$ lungs. Scale bar, A, 200 μm, B, 100 μm, C, 50 μm.

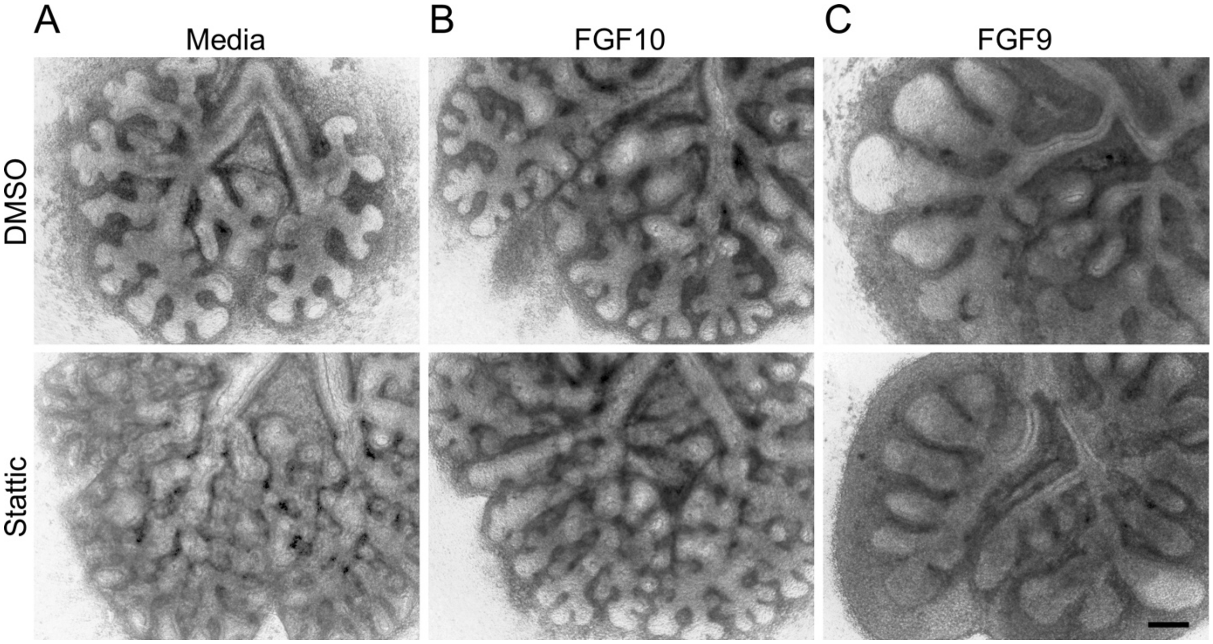


**Fig. S6. Inhibition of STAT3 has minimal effects on FGF9 and FGF10 signaling.**

E11.5 wild-type lung explants were treated with vehicle (DMSO) or the STAT3 inhibitor, Stattic, and simultaneously treated with (A) media, or media containing (B) FGF10 or (C) FGF9. Explants were photographed after 48 h. Images representative of at least 3 embryos. Scale bar, 200 $\mu$m.
